# Interferon-inducible miR-128 modulates HIV-1 replication by targeting TNPO3 mRNA

**DOI:** 10.1101/195511

**Authors:** Aurore Bochnakian, Dimitrios G Zisoulis, Adam Idica, Anjie Zhen, Vineet N KewalRamani, Iben Daugaard, Matthias Hamdorf, Scott Kitchen, KyeongEun Lee, Irene Munk Pedersen

**Affiliations:** Department of Molecular Biology and Biochemistry, Francisco J. Ayala School of Biological Sciences, University of California - Irvine, Irvine, CA 92697-3900; UCLA AIDS Institute, Los Angeles, CA 90095; Basic Research Laboratory, Center for Cancer Research, National Cancer Institute, Frederick, MD 21702-1201; Department of Biomedicine, Aarhus University, DK-8000 Aarhus C, Denmark

**Keywords:** miR-128, HIV-1, TNPO3, nuclear import, restriction factor

## Abstract

The HIV/AIDS pandemic remains an important threat to human health. We have recently demonstrated that a novel microRNA (miR-128) represses retrotransposon (LINE-1 or L1) by a dual mechanism, by directly targeting the coding region of the L1 RNA and by repressing a required nuclear import factor (TNPO1). We have further determined that miR-128 represses the expression of all three isoforms of TNPO proteins (transportins, TNPO1,-2 and TNPO3). Here, we establish that miR-128 also controls HIV-1 replication by repressing TNPO3. TNPO3 is well established to regulate HIV-1 nuclear import and viral replication. Here, we report that the type I interferon inducible miR-128 directly targets two sites in the TNPO3 mRNA, significantly down-regulating TNPO3 mRNA and protein expression levels. Manipulation of miR-128 levels in HIV target cell lines and in primary human CD4 T-cells by over-expression or knockdown showed that modulation of TNPO3 by miR-128 affects HIV-1 replication but not MLV infection. In addition, we found that miR-128 modulation of HIV-1 replication is reduced with TNPO3-independent HIV-1 virus and in cells depleted of CPSF6, suggesting that miR-128-indued TNPO3 repression is partly required for miR-128-induced inhibition of HIV-1 replication. Finally, challenging miR-modulated Jurkat cells or primary CD4 T-cells with wildtype, replication-competent HIV-1 shows that miR-128 significantly delays spreading infection. Thus, we have established a novel role of miR-128 in anti-viral defense in human cells, inhibiting HIV-1 replication partly by targeting TNPO3.

## IMPORTANCE

HIV-1 is the causative agent of AIDS. During HIV-1 infection, type I interferons (IFNs) are induced and their effectors limit HIV-1 replication at multiple steps in its life cycle. However, the underpinning mechanisms of INFs are still largely unknown. In this study we identified the interferon-inducible miR-128, as a novel antiviral mediator, which suppresses the expression of the host gene TNPO3 required for HIV-1 replication. Specifically, TNPO3 disruption (by miR-128) leads to accumulation of CPSF6 in the cytoplasm and inhibition of viral nuclear entry and viral replication. Finally, the miR itself or the cellular pathway identified as modulating virus replication may prove to be novel candidates for the development of therapeutic interventions.

## INTRODUCTION

### Human immunodeficiency virus (HIV) infection

The HIV/AIDS pandemic remains an important threat to human health. To date more than 34 million people are infected with HIV. Unique characteristics make HIV-1 difficult to eradicate. HIV-1 uses and evades the immune system, replicating in and destroying immune cells during active replication. To develop novel and widely applicable strategies, a thorough understanding of the cellular determinants of HIV-1 replication and viral latency is crucial, including epigenetic changes and transcriptional and post-translational regulation (1-7). Due to its limited genome (10 kb) HIV-1 has to rely on cellular co-factors to progress through its lifecycle. Host factors are not only needed for productive HIV-1 replication but also for latency establishment and reactivation whereas restriction factors protect the cells against infection and provide innate immunity (8-14).

### miRs and their role as viral restriction factors

microRNA (miRNA or miR) biogenesis begins in the nucleus with transcription, to create the long primary miR (pri-miR) including a hairpin that contains the mature sequence. The hairpin is excised by the microprocessor that includes Drosha, an RNase III enzyme, and its cofactor DGCR8, producing the 60–70 nucleotide precursor miRs (pre-miR). The pre-miR is exported out of the nucleus by Exportin-5 where another RNase III enzyme, Dicer, processes the pre-miR into the 21–23 nucleotide duplex miR. The strand destined to be the mature sequence is then loaded onto Argonaute (Ago), forming the miR-induced silencing complex (miRISC) along with other proteins (15,16). Using imperfect base pairing, miRs guide RISC to specific mRNAs to down-regulate their expression by triggering mRNA destabilization or translational repression (17,18). miR-target recognition is often dependent on a short, six nucleotide *seed sites*, which perfectly complement the 5’ of the miR (position 2-7) (19). Both miRs and the mRNA binding sites are highly conserved (19,20). miRs exemplify the emerging view that non-coding RNAs may equal proteins in regulatory importance. The majority of the human transcriptome is predicted to be under miR regulation, positioning this post-transcriptional control pathway within every major genetic cascade (21). miRs can regulate and coordinate development, modulate self-renewal, differentiation and self fate establishment (12-25). In addition, Dicer and Drosha knock-out experiments has demonstrated that miR pathways repress HIV-1 replication and contribute to the maintenance of latency (26). Furthermore, profiling studies suggests that miRs in general are down-regulated upon T-cell activation, suggesting increased target de-repression during optimal HIV-1 infection (27-29). Also, miRs have been described to function in complex ways in the intersection between host and pathogen for example as an anti-viral defense mechanism and/or as a facilitator of latency (30,31). miRs has been proposed to negatively affect HIV-1 by directly targeting of the viral RNA genome and or by repressing virus-dependent cellular co-factors, these miRs include miR-29, -150,-28, -125b, 223 and 382 (32). The complex interplay between miRs and their mRNA targets during HIV-1 infection, replication and latency needs further investigations.

Work in colleagues and our laboratory established that interferon-induced miRs represse viral replication of Hepatitis C Virus (HCV) (30,33-35). As the life cycle of long-interspaced-elements-1 (L1) mimics that of a virus, we next asked the question whether miRs also regulate L1 activity. L1 retrotransponsons and related SINE elements make up approximately 35% of the human genome, with some L1 elements still being active, regardless of multiple cellular restriction mechanisms (36-41). We recently performed an anti-miR lentiviral library screen to identify miRs involved in the regulation of *de novo* L1 retrotransposition. miR-128 was identified as a negative regulator of L1 and further characterization led to the discovery of a novel principle by which miR-128 represses *de novo* retrotransposition of L1 elements in somatic cells, including cancer cells, cancer stem cells and iPSCs, by directly binding L1 RNA and targeting it for degradation (42). This finding was surprising as the most well understood mechanism by which miRs function relies on the binding and regulation by one miR to multiple cellular mRNAs, whose protein products typically function in a specific pathway within the cell. We therefore explored if miR-128 might also regulate cellular proteins involved in L1 retrotransposition. We used bioinformatics analysis in combination with miR-128 RNA immune-purification (RIP) RNA Smart sequencing approaches and identified the mRNA for a nuclear import factor (TNPO1) as a target of miR-128 binding. Our recent results (*manuscript under revision*) determines that TNPO1 is a functional target for miR-128, by directly targeting TNPO1 for degradation, which result in the L1 ribonucleoprotein (RNP) complex being trapped in the cytoplasm and L1 retrotransposition and genomic integration repressed. Thus miR-128 represses L1 retrotransposition by two independent mechanisms: 1) repression of a key co-factor required by L1 (TNPO1), and 2) directly binding L1 RNA (42). The TNPO family of proteins contains, TNPO1, -2 and TNPO3. Interestingly miR-128 has predicted binding sites in all three TNPO protein isoforms.

Here we report that the type I interferon induced microRNA miR-128 regulates TNPO3 mRNA and protein expression levels in cell lines (HeLa, Jurkat, and THP-1 cells) and in primary CD4 T-cells, by directly targeting TNPO3 mRNA. Infection studies using wild-type and TNPO3-independent N74D mutant HIV-1 reporter virus demonstrate that miR-128 significantly represses HIV-1 replication in cell lines and in CD4 T-cells, and that miR-128-induced HIV-1 repression is partly dependent on TNPO3 reduction (N74D HIV-1 showing de-repression). The cleavage and polyadenylation specificity factor subunit 6 (CPSF6), one of SR proteins, is a conditional restriction factor for HIV-1, and can be transported to the nucleus by TNPO3, the nuclear importer of SR proteins. When TNPO3 is removed from cells, CPSF6 accumulates in the cytoplasm and restricts HIV-1 infection (51). CPSF6-depletion studies show a partial rescue of HIV-1 inhibition caused by TNPO3 reduction. Finally, infection studies using replication competent HIV-1_NL4-3_ virus a significant delay in viral spreading in miR-128 overexpressing Jurkat and primary CD4+ T-cells. Together these studies support the idea that when TNPO3 is disrupted (by miR-128), then CPSF6 accumulates in the cytoplasm and inhibits HIV-1 replication.

## RESULTS

### miR-128 regulates TNPO3 expression levels

Three in the importin gene family, TNPO1, TNPO2 and TNPO3, contain predicted miR-128 binding sites (Supplemental Figure S1). This is noteworthy, as TNPO3 is required for successful nuclear import of the HIV-1 pre-integration complex (PIC) and viral replication (43-49). As mentioned, CPSF6 is a conditional HIV-1 co-factor, aiding in nuclear steps of virus replication in cells retaining TNPO3 function (48). If TNPO3 is disrupted, CPSF6 accumulates in the cytoplasm and inhibits HIV-1 viral replication (49-53). Of note, we and others have determined that type I interferon induces miR-128 levels, suggesting a role for miR-128 in anti-viral defense in human cells (Supplemental Figure S2 and (30,54)). Furthermore, miR-128 is expressed in HIV-1 target cells including CD4 T-cells and blood-derived monocytes (Supplemental Figure S2 and (54-58)). Thus, if miR-128 represses expression of TNPO proteins, including TNPO3, we would predict that miR-128 represses HIV-1 replication.

To explore this possibility, we generated stably transduced miR-modulated cells lines (miR-128 overexpressing, anti-miR-128 in which endogenous miR-128 is neutralized and miR controls) of HeLa cells. We validated that miR-128 expression levels was modulated as expected and at physiological relevant levels similar to changes induced by cytokines and growth factors (30,54) by miR specific qPCR (miR-128 was induced 4-5 fold in miR-128 overexpressing cells, while miR-128 was reduced by 40-50% in anti-miR-128 expressing HeLa cells) (Supplemental Figure S3).

Next, we evaluated TNPO3 levels in cell lines in which miR-128 was reduced or overexpressed. The level of TNPO3 mRNA was significantly reduced in miR-128 overexpressing HeLa cells compared to miR controls, whereas TNPO3 mRNA levels were increased when miR-128 was neutralized by anti-miR-128 (Figure 1A, left panel). To rule out the possibility that miR-128 overexpression using a lentiviral delivery strategy itself caused reduction in TNPO3 mRNA levels, we next examined the levels of TNPO3 mRNA in HeLa cells after transiently transfecting with miR-128, anti-miR-128 or control miR oligonucleotides (miR mimics and anti-miRs). 48 hours after transfection, significantly enhanced expression of TNPO3 mRNA was observed in HeLa cells transfected with anti-miR-128 oligonucleotide and significant reduction of TNPO3 mRNA was observed in HeLa cells transfected with the synthetic miR-128 mimic (Supplemental Figure S4). To exclude the possibility that the observed miR-128 effect was cell-type specific (limited to HeLa cells), we next generated stable miR-modulated THP-1 cell lines and determined that miR-128 significantly represses TNPO3 mRNA expression levels, in contrast to anti-miR-128 which significantly enhanced TNPO3 mRNA expression levels in THP-1 cells relative to miR controls (Figure 1A, middle panel). Substantial changes in TNPO3 mRNA levels were also observed by transient transfections of miR mimics in Jurkat and THP-1 cells (Supplemental Figure S4). Next, we wished to evaluate the effect of miR-128 on TNPO3 expression in primary CD4+ cells. We obtained PBMCs from healthy donors and isolated CD4-positve T-cells by Ficoll and negative MACS (Magnetic-activated cell sorting separation). Primary CD4+ cells were transiently transfected with miR mimic oligonucleotides (as described above), RNA was isolated after 48hrs and TNPO3 mRNA expression levels were determined by qPCR analysis. miR-128 significantly decreased the expression of TNPO3 mRNA in CD4+ T cells relative to miR control and anti-miR-128 showed the same significant phenotype that we observed in the other cell types (Figure 1A, right panel). Finally, we examined TNPO3 protein expression by western blot analysis and conformed that miR-128 decreased the protein expression of TNPO3 which correlates with the levels of TNPO3 mRNA levels in HeLa and CD4+ T cells overexpressing miR-128 (Figure 1B, right panels, quantification, lower panel). These combined results suggested that miR-128 regulates TNPO3 expression levels in multiple cell types including primary HIV-1 target cell type, CD4+ positive T-cells.

**Figure 1:**
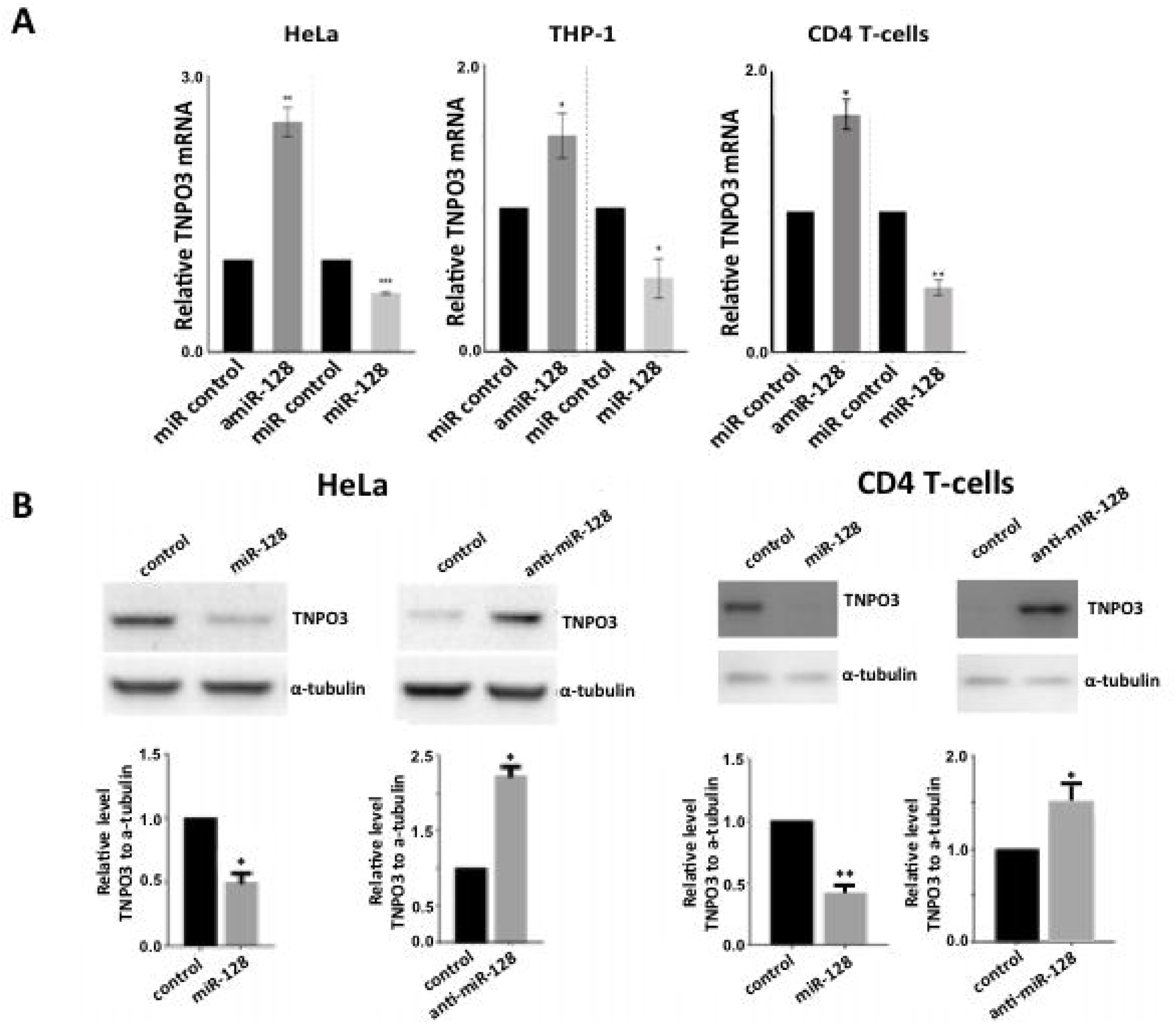
Identification and verification of TNPO3 as a cellular target of miR-128. **(A)** Relative levels of TNPO3 RNA to B2M in HeLa or THP-1 cells stably transduced or primary CD4+ T-cells transiently transfected with control miR, anti-miR-128 or miR-128 are shown as mean ± SEM (n=3, independent biological replicate) **(B)** Western blot analysis of TNPO3 and alpha-tubulin protein levels in lysates from HeLa cells stably transduced with control, anti-miR-128 or miR-128 lentiviral constructs or primary CD4+ T-cell transiently transfected with miR mimics (as described above in (A)). (Shown is one representative out of 3 independent biological replicates. Quantification of n=3). Throughout figure, *P<0.05; **P<0.01 ***, p<0.001 by two-tailed Student’s *t* test.

### miR-128 directly interacts with TNPO3 mRNA

Next, we explored the mechanism by which miR-128 regulates TNPO3 levels. By performing bioinformatics analysis (using TargetScan and miRBase 59,60) we identified two predicted miR-128 binding sites in TNPO3 mRNA, one 7-mer seed site in the coding region sequence (CRS) of TNPO3 mRNA (Site #1) and a second 7-mer seed site in the 3’UTR of TNPO3 mRNA (Site #2) (see cartoon Figure 2A, top). TNPO3 CRS and 3’UTR sequences harboring the two potential miR-128 binding sites were cloned into a dual-luciferase reporter constructs (pEZX-MT05). In addition, a perfect 23 nt miR-128 binding sequence (positive control) luciferase construct was generated. HeLa cells were transfected with one of the three miR-128 binding site-encoding plasmids (Site #1 CDS TNPO3, Site #2 3’UTR TNPO3 or miR-128 perfect binding site) in addition to mature miR-128 or miR control oligonucleotide mimics (see Figure 2A). To determine if the specific miR-128 seed sequences are responsible for the interaction with miR-128, we also measured luciferase activity in HeLa cells transfected with mature miR-128 (or control miR mimics) along with either CRS or 3’UTR TNPO3 binding sites in which mutations had been introduced (see Figure 2A). While luciferase activity was significantly reduced in cells transfected with miR-128 and encoding the WT binding sites (Site #1 and Site #2) of TNPO3 as well as with the positive control (Figure 2B), the decrease in luciferase activity in HeLa cells transfected with mutant TNPO3 and miR-128 were not significant although it was not completely restored to control levels (Figure 2C). These results indicate that miR-128 targets TNPO3 mRNA and both predicted seed sites are responsible for their interaction (Figure 2C).

**Figure 2:**
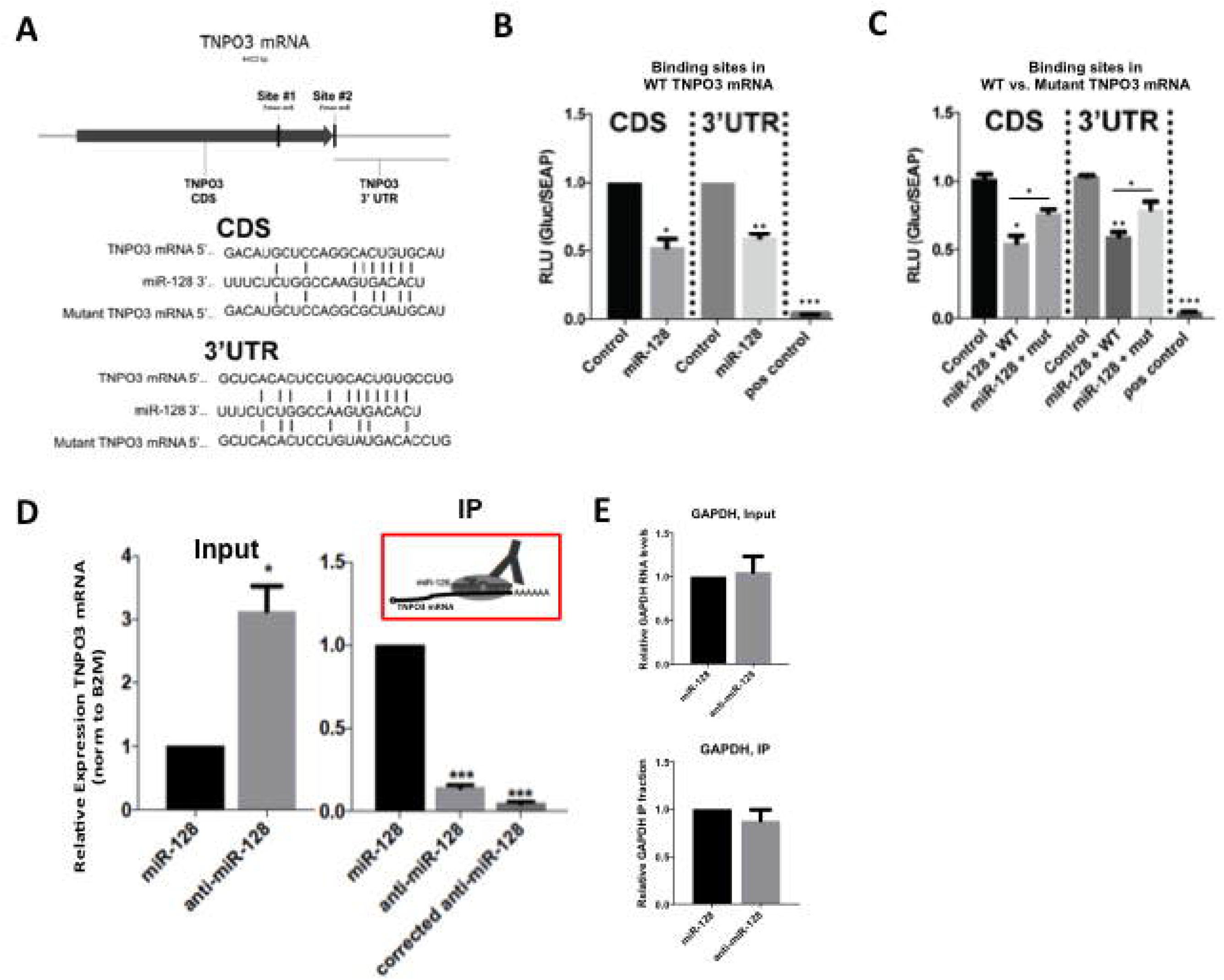
miR-128 represses TNPO3 by binding directly to two sites in TNPO3 mRNA. **(A)** Schematic of the two predicted miR-128 binding sites in TNPO3 mRNA, which includes a 7-mer seed site (Site#1) in the coding reading sequence (CRS) and 7-mer seed site (Site#2) in the 3’UTR of TNPO3 mRNA (top panel). The predicted base pairing of miR-128 to the seed sequence of the two wild-type (WT) binding sites in TNPO3 mRNA as well as a representation of mutations, which we generated in the seed sequence (mutant) are shown (bottom panels). **(B)** Relative luciferase activity in HeLa cells transfected with plasmids expressing a Gaussia luciferase gene fused to the one of the two predicted wild-type (WT) binding sites in TNPO3 or a positive control sequence corresponding to the 22 nucleotide perfect match of miR-128 and co-transfected with control or mature miR-128 mimics were determined 48 hours post-transfection (n=3). **(C)** Relative luciferase activity in HeLa cells transfected with plasmids expressing the luciferase gene fused to the WT or either mutated binding site (mutant) and co-transfected with control or mature miR-128 mimics were determined 48 hours post-transfection (n=3). **(D)** Argonaute-RNA immuno-purification in HeLa cell lines stably transduced with miR-128 overexpression or miR-128 neutralization (anti-miR-128) was performed. Relative amounts of TNPO3 mRNA normalized to B2M was determined for input samples (left panel “input – TNPO3”, n=3). Relative fraction of TNPO3 transcript amounts associated with immune-purified Ago complexes is shown for IP samples, TNPO3 fractions normalized to the amount of TNPO3 in input are shown as “corrected” (top right panel “IP – TNPO3”, n=3). **(E)** Relative amount of GAPDH in the same input and IP samples were determined as a negative control (n=3). Throughout figure, n=3 independent biological replicates, mean **±** SEM *P<0.05; **P<0.01; ***, p<0.001 by two-tailed Student’s *t* test.

Furthermore, Argonaute (Ago) complexes containing miRs and target mRNAs were isolated by immuno-purification and assessed for relative complex occupancy by the TNPO3 mRNA to determine if miR-128 directly binds TNPO3 mRNA in HeLa cells (see cartoon in Figure 2D), as previously described (42,61). The relative level of TNPO3 mRNA was significantly lower in cells stably overexpressing miR-128 when compared to those expressing anti-miR-128 constructs, as expected (Figure 2D, Input, left panel). When correcting for the lower expression level of TNPO3 mRNA, (because of lower miR-128 expression levels), which may underestimate the scale of the effect, the relative fraction of immunopurified-Ago-bound TNPO3 mRNA significantly increased when miR-128 was overexpressed, compared to anti-miR-128 control (Figure 2D, IP, right panel). In contrast, miR-128 did not repress GAPDH mRNA expression levels or immuno-purified GAPHD mRNA, as expected (Figure 2E). This result suggests that high levels of miR-128 lead to higher levels of TNPO3 mRNA being bound in Ago-complex and TNPO3 expression is regulated directly by miR-128. These data imply that miR-128 represses TNPO3 expression via a direct interaction with the target sites located in the CRS and 3’UTR of the TNPO3 mRNA.

### miR-128 represses HIV-1 replication

In order to study if type I interferon-induced miR-128 functions as a novel anti-viral mediator during HIV-1 infection in human cells and is specific to restricting HIV-1 replication, we employed VSV-G pseudotyped HIV-1 vector that contains either luciferase (NLdELuc) or red fluorescent protein gene (HIV-1-RFP) as a reporter gene. HIV-1 env has been deleted in these vectors and the nef gene has been replaced with a reporter gene [49]. miR modulated Jurkat cells expressing miR control or miR-128 were infected with either HIV-1-RFP/VSV-G or MLV-RFP/VSV-G. Then their infection was examined by FACS analysis for RFP expression after 48hrs. These experiments showed that miR-128 significantly inhibits HIV-1-RFP/VSV-G, but did not substantially block MLV-RFP/VSV-G in Jurkat cells (Figure 3A and 3B).

**Figure 3:**
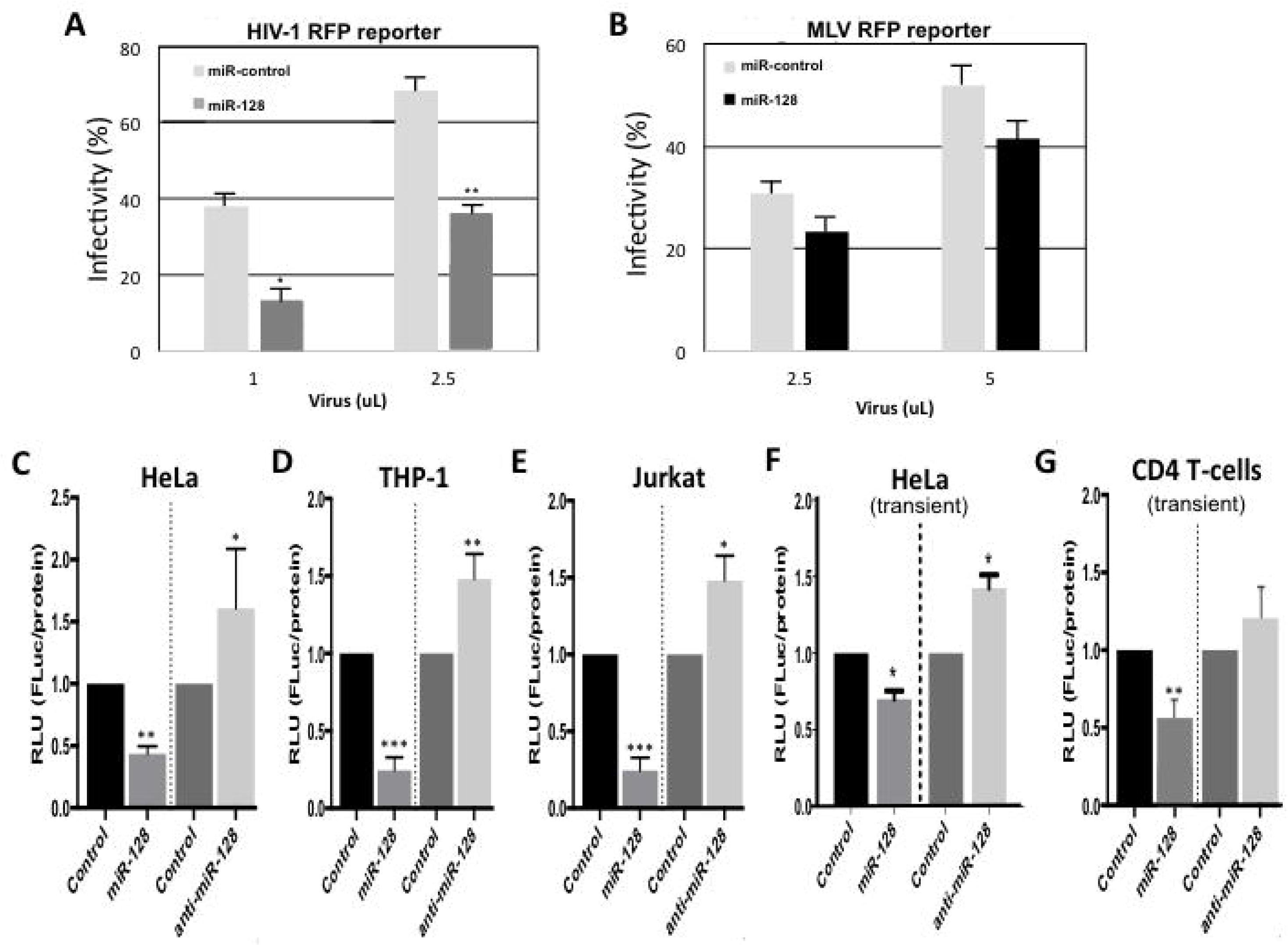
miR-128 inhibits HIV-1 replication of a single-cycle HIV-1 reporter virus. **(A and B)** Stable miR-modulated Jurkat cells (miR control or miR-128 overexpressing) were spinfected with VSV-G pseudotyped HIV-1 or MLV vectors that contains red fluorescent protein gene (HIV-1-RFP or MLV-RFP/VSV-G). The fraction of infected cells was measured by FACS for RFP expression after 48hrs and equal infectivity between samples were verified by quantification of RFP expression. **(C-E)** miR-modulated HeLa, THP-1, or Jurkat cells (miR-128, anti-miR-128 or control miR) were generated by stable transduction. Alternatively HeLa cells **(F)** or primary human CD4+ T-cells **(G)**, isolated from healthy blood donors by Ficoll and negative MACS separation, were transiently manipulated. CD4+ T-cells were activated for 2 days with CD3/CD28 and cultured in the presence of 30U/ml IL-2. Activated and IL-2 stimulated CD4+ T-cells or HeLa cells were then transiently transfected with miR mimics (miR-128, anti-miR-128 or controls miR oligonucleotides). Both stable and transient miR modulated cells were then spinfected with the HIV-1 reporter virus as described in (A and B). After 48 hours luciferase was measured. Throughout figure, n=3 independent biological replicates, mean ± SEM *P<0.05; **P<0.01; ***, p<0.001 by two-tailed Student’s *t* test.

Next, we wished to examine the specificity of the miR-128 effect, by comparing miR-128 overexpression versus miR-128 depletion (by anti-miR-128) to miR control expressing cells. We evaluated the effect of miR-128 in three different cell lines by generating stable miR-128-modulated HeLa, Jurkat (T-cell line), and THP-1 cells (monocytic cell line). Cells were infected with HIV-1 reporter virus pseudotyped with VSV-G by spinoculation, and infection monitored by measuring luciferase activity 48hrs after infection. We found that miR-128 significantly inhibited HIV-1 infection whereas anti-miR-128 significantly enhanced HIV-1 infection, relative to miR controls in HeLa, Jurkat and THP-1 cells (Figure 3C-E). To rule out the possibility that the miR effect was an artifact of lentiviral genomic integration, we tested transient miR modulated HeLa cells using miR/anti-miR mimic oligonucleotides. We observed that transient miR-128 overexpression significantly inhibited HIV-1 infection and miR-128 neutralization enhanced HIV-1 replication relative to miR controls. Not surprisingly, the transient miR effect was less significant relative to stable miR transductions, as only 60-80% of cells are transfected with miR (data not shown) (Figure 3F). Next, primary human CD4+ T-cells were tested. The isolated CD4+ T-cells were stimulated with 30 IU/ml IL-2, activated with CD3/CD28 for 2 days, and transiently transfected with miR mimics (miR-128, anti-miR-128 and control miR oligonucleotides) for 48hrs. These cells were then infected with HIV-1 reporter virus by spinoculation and luciferase activity was measured after 48hrs. miR-128 significantly inhibited HIV-1 infection in primary CD4+ T-cells, to a similar extent as observed with transient miR modulated of HeLa cells (Figure 3G). anti-miR-128 did not affect replication of the HIV-1 infection in any substantial fashion. This combined data confirmed that miR-128 specifically and significantly inhibits infection of a single cycle HIV-1 infection in cell lines (HeLa, Jurkat and THP-1 cells) and in HIV-1 target cells, CD4+ T-cells.

### miR-128-induced HIV-1 inhibition is partly dependent on TNPO3

Next, we wished to explore whether miR-128 induced-TNPO3 repression is required for HIV-1 repression. We took advantage of the N74D CA mutant single-cycle HIV-1 reporter virus pseudotyped with VSV-G (N74D NLdELuc). The single mutation at N74 in HIV-1 capsid has been shown to affect the sensitivity of HIV-1 infection to depletion of various nuclear import factors including TNPO3 (49,62,63).

In order to dissect the role of miR-128-induced TNPO3 repression, we generated miR-modulated and TNPO3-modulated (shTNPO3, TNPO3 overexpressing or plasmid controls) HeLa cell lines. All cell lines were validated for TNPO3- and miR-128-expression levels (Figure 4A top right, Figure 2B and Supplemental Figure S3). miR-128 and TNPO3-modulated cells were spinoculated with WT or N74D mutant HIV-1 reporter virus. As shown before (Figure 3C), miR-128 significantly reduced WT HIV-1 replication and anti-miR-128 enhanced WT HIV-1 replication, relative to miR control HeLa cell lines (Figure 4A). TNPO3 stable knockdown HeLa cells by shRNA transduction (shTNPO3) showed significant HIV-1 inhibition and overexpression of TNPO3 expression increased WT HIV-1 infection relative to control cells, as expected (Figure 4A). When comparing WT HIV-1 with N74D HIV-1 infection in these cells known to be insensitive to TNPO3 knockdown, we observed that neither TNPO3 knockdown or overexpression significantly affected N74D HIV-1 infection as previously shown by other groups (Figure 4B, right panel) and we verified that the N74D CA mutant HIV-1 is not dependent on TNPO3 for nuclear entry and viral replication. The effect of miR-128-induced inhibition of HIV-1 viral replication, was significantly reduced in N74D infected HeLa cells, as compared to WT HIV-1 reporter replication, relative to miR control HeLa cell lines. However the potency of inhibitory effect of N74D HIV-1 by miR-128 was significantly less relative to the effect on WT HIV-1 reporter virus (Figure 4B, left panel).

**Figure 4:**
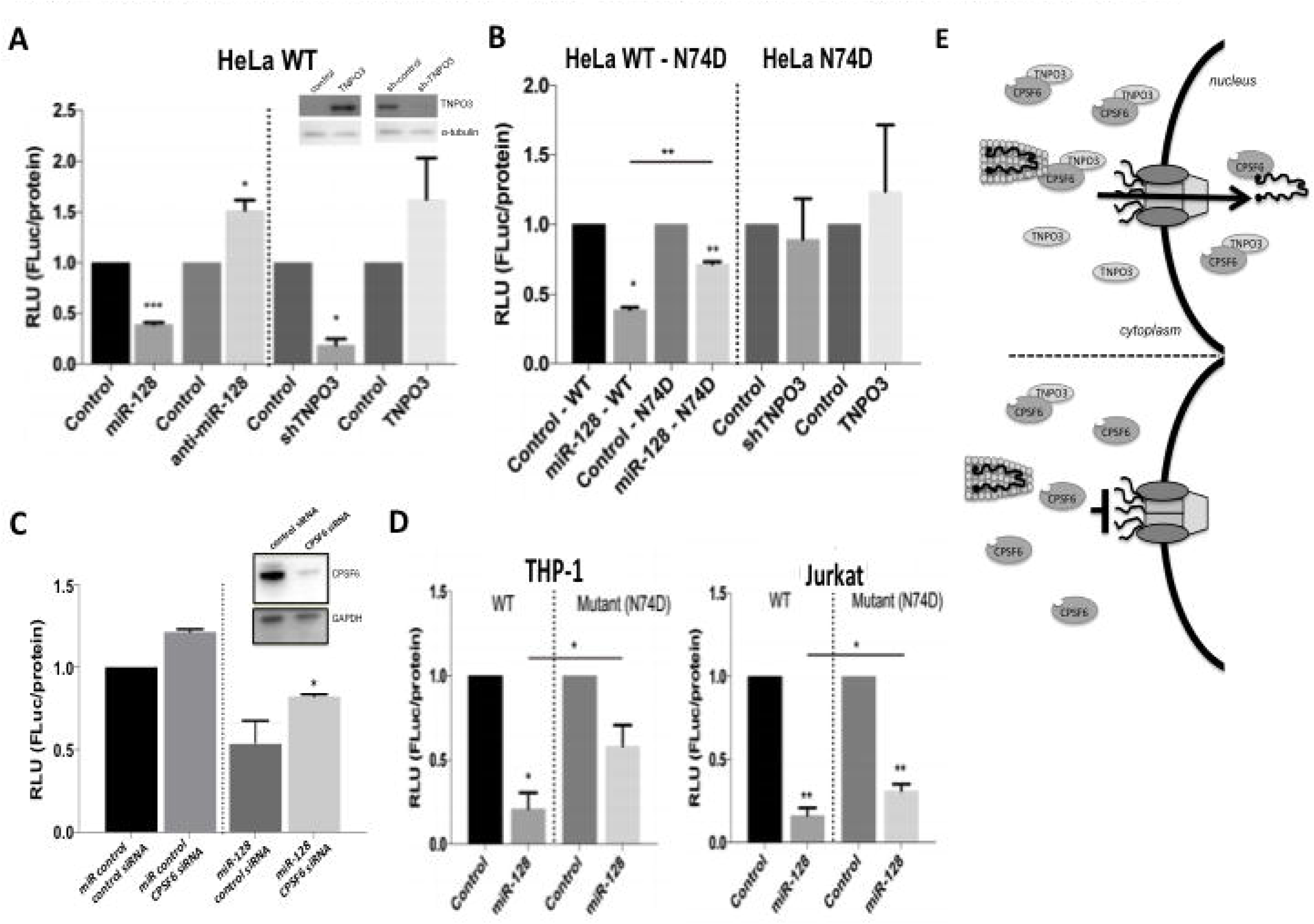
miR-128-induced HIV-1 repression is partly dependent on TNPO3. **(A)** miR-modulated HeLa cells (miR-128, anti-miR-128 or control miR) or TNPO3-modulated HeLa cells (shTNPO3, TNPO3-overexpressing or plasmid controls) were generated by stable transduction, spinfected with luciferase encoding VSV-G-pseudotyped RFP HIV-1 reporter virus and luciferase were measured after 48 hours. Equal infection was determined by counting RFP positive cells (n=3). Modulation of TNPO3 protein expression in HeLa cells was verified by western blot analysis of TNPO3 and alpha-tubulin (top right). One representative example is shown out of three. **(B)** miR-modulated HeLa cells or TNPO3 modulated HeLa cells were spinfected with the wildtype (WT) or the N74D variant (TNPO3 independent) of the VSV-G-pseudotyped RFP HIV-1 reporter virus and luciferase were measured after 48 hours. **(C)** miR control or miR-128 overexpressing HeLa cells were transfected with CPSF6 or control siRNA oligonucleotides. Modulation of CPSF6 protein expression was verified by western blot analysis of CPSF6 and alpha-tubulin (top right). Cells were spinfected with RFP HIV-1 reporter virus and luciferase were measured after 48 hours. Equal infection was determined by counting RFP positive cells. **(D)** miR-modulated THP-1 and Jurkat cells (miR-128, anti-miR-128 or control miR) were generated by stable transduction, and spinfected with either the WT luciferase encoding VSV-G-pseudotyped RFP HIV-1 reporter virus or the N74D mutant HIV-1 reporter virus. Luciferase was measured after 48 hours. Throughout figure, n=3 independent biological replicates, mean ± SEM *P<0.05; **P<0.01; ***, p<0.001 by two-tailed Student’s *t* test. **(E)** Model of TNPO3/CPSF6’s effect on HIV-1 replication. CPSF6 functions as a conditional inhibiting factor, acting with low levels of TNPO3. If TNPO3 is disrupted (for example by miR-128), CPSF6 accumulates in the cytoplasm and impairs viral nuclear entry and viral replication of HIV-1 (as shown in the bottom panel of cartoon).

The cellular factor CPSF6 mostly localizes in the nucleus. Interestingly, it was shown to interact with HIV-1 CA (capsid) protein (but not the N74D mutant), and CPSF6 becomes anti-viral when it localizes in the cytoplasm (49). As explained, when TNPO3, a carrier of SR proteins to the nucleus, is removed from cells, CPSF6 accumulates in the cytoplasm and blocks HIV-1 infection at the nuclear entry process by sequestering HIV-1 PIC associated CA (49-52) (and see Figure 4E). Therefore, we next tested if miR-128-induced TNPO3 repression causes the increase of cytoplasmic CPSF6, and specifically whether miR-128 repression of HIV-1 infection also requires CPSF6. miR-modulated HeLa cells were challenged with HIV-1 reporter virus after transient transfection with siRNA targeting CPSF6 or a non-specific control siRNA. CPSF6 knockdown was validated by western blot (Figure 4C, top right). miR-128-induced inhibition of HIV-1 replication was partly restored in CPSF6-depleted HeLa cells, relative to miR-128 overexpressing cells challenged with control siRNAs (Figure 4C).

Finally we evaluated the anti-viral effect of miR-128 on WT and N74D mutant HIV-1 reporter virus using the more relevant cell lines such as THP-1 or Jurkat cells. We observed that miR-128 induced significant inhibition of WT HIV-1 infection in THP-1 cells (Figure 4D) and Jurkat (Figure 4E) as previously established (Figure 3D and 3E). When challenging miR-modulated Jurkat and THP-1 cells with the N74D mutant reporter virus, we found that miR-128-induced inhibition of N74D viral replication was significantly de-repressed as compared to repression of WT viral replication. However, miR-128 still significantly reduced N74D HIV-1 reporter activity, as compared to miR control cell lines (Figure 4D and 4E). These studies suggest that miR-128-induced inhibition of HIV-1 viral replication (of the HIV-1 reporter virus) is partly dependent on reduction of TNPO3 expression in target cells.

### miR-128 delay viral infection and replication of HIV-1

Next we wished to determine if miR-128-induced repression of TNPO3 causes accumulative anti-viral effects on HIV-1 viral replication, as compared to analysis of the single-cycle HIV-1 reporter virus. For this reason we performed viral infection of miR-modulated Jurkat cells, using replication competent WT HIV-1 virus (HIV-1_NL4-3_). miR-modulated Jurkat cells were infected with equal amounts of WT virus verified by normalizing to RT Units. We determined that miR-128 significantly reduced viral spreading of wildtype HIV-1 both in conditions of low and high dose HIV-1, as determined by p24 assays, as compared to Jurkat cells transduced with control miR (Figure 5A). Viral spreading was delayed by 4 days by miR-128, and viral load was reduced to levels where miR-128 cells, unlike miR control cells, maintained viability during viral replication when challenging cells with high dose HIV-1 (Figure 5A).

**Figure 5:**
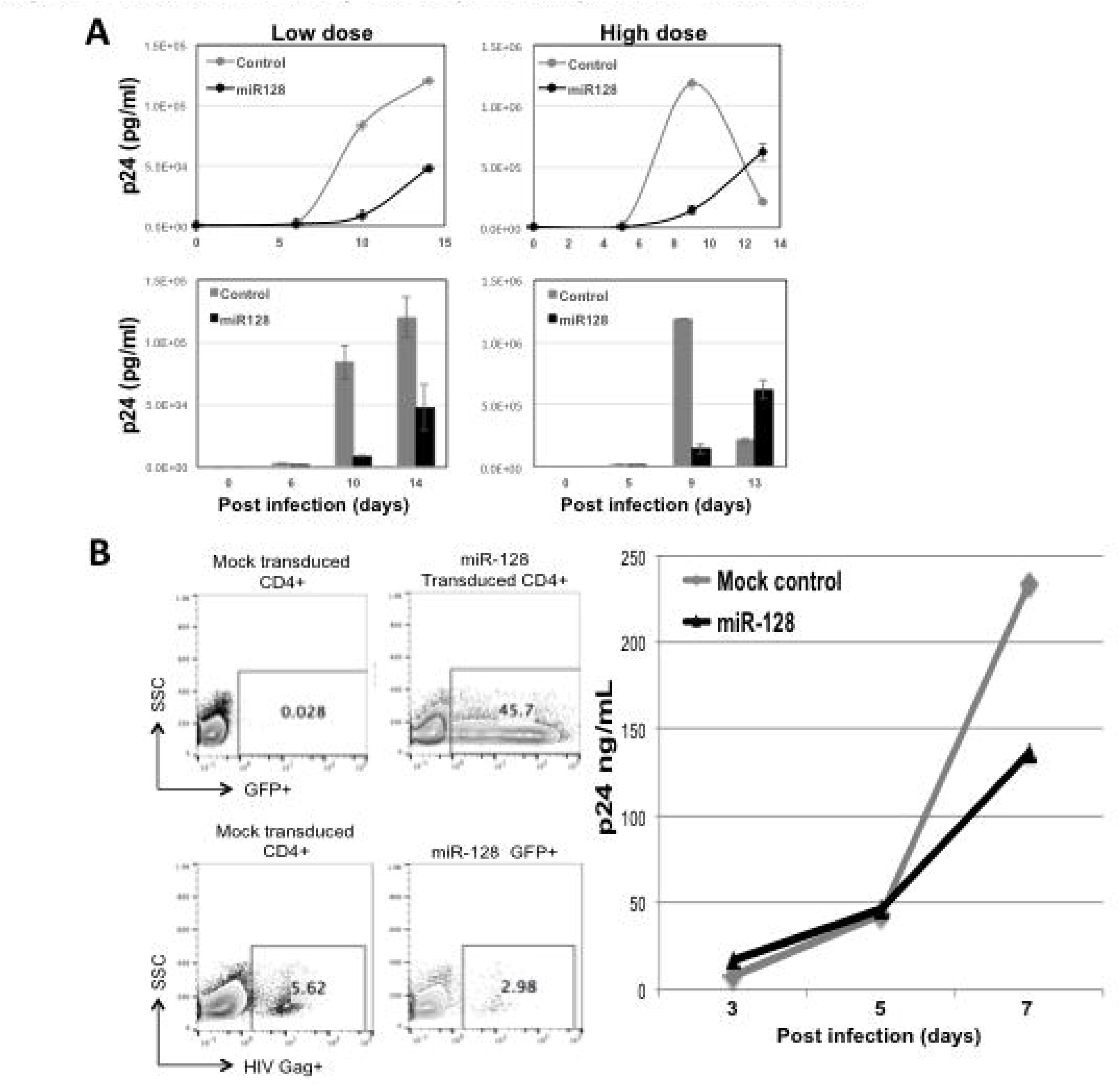
miR-128 delays viral spreading of wild-type HIV-1. **(A)** miR-modulated Jurkat cells (miR-128 or control miR) were generated by stable transduction, and spinfected with wildtype, replication competent HIV-1 virus WT_NL4-3_ and spreading was determined by collecting supernatant for p24 assays analysis. Equal infectivity was determined by measuring RT units. (n=3 independent biological replicates). **(B)** CD4+ cells were sorted from healthy donors PBMCs, activated with anti-CD3/anti-CD28 for 2 days and transduced with miR lentivirus. After 2 days cells were infected with HIV-1 NL4-3 (100ng/10E6) for 7 days. Supernatant were collected on day 3,5 and 7 for p24 assays. On day 7 infected cells were harvested and stained for intracellular anti-gag. Level of transduction infection as presented by percent gag+ shown among GFP-positive transduced cells. miR-128 transduced CD4+ T-cells was characterized by a delay in viral infection, relative to mock CD4+ T-cell controls. (n=3 technical replicates of 2 independent biological replicates).

Finally, we evaluated the anti-viral effect of miR-128 on primary CD4+ T-cells. CD4+ T cells were isolated from healthy donors PBMCs and sorted by flow cytometry, cells were activated with anti-CD3/anti-CD28 for 2 days and transduced with miR lentivirus. After 2 days, cells were infected with HIV-1_NL4-3_ (100ng/10E6) for 7 days. Supernatant were collected on day 3, 5, and 7 for p24 assays. On day 7 infected cells were harvested and stained for intracellular anti-gag. Level of infection is presented as percent gag+ shown among GFP-positive transduced cells as previously described (64,65). miR-128 transduced CD4+ T-cells was characterized by a delay in viral infection, relative to mock controls. Despite the inefficiency of transient transfection in the primary cells, CD4+ T cells expressing miR-128 showed increased resistance to HIV-1 replication over the first week of infection, comparable to what was observed in the Jurkat cell lines. Unfortunately, the viability of the primary stimulated cells was limited such that the infection time course was shorter than the experiment in the Jurkat lines.

In summary, these experiments support the idea that miR-128 significantly repress viral replication and delay viral spreading of HIV-1, and that the miR-128 asserted anti-viral defense mechanism is partly dependent on targeting TNPO3.

## DISCUSSION

Our present data provides the first evidence that the small RNA (miR-128) functions as an anti-viral mediator restricting HIV-1 replication and viral spreading. The finding that miR-128 can be induced by type I interferon in HIV-1 target cells (CD4+ T-cells and blood-derived monocytes), supports the idea that miR-128 functions as a novel host restriction factor. However, additional studies are required to decipher the complex interplay between miR-128 and different classes and subclasses of interferon, including whether inhibition of miR-128 can release IFN restriction on HIV-1 infection.

Mechanistically, we demonstrate that miR-128 reduces TNPO3 mRNA and protein levels in cell lines (HeLa, Jurkat and THP-1 cells) and in primary CD4+ T-cells, by directly targeting two sites in the TNPO3 mRNA, one in the 3’UTR and one in the coding region sequence (CRS). Analysis of miR binding sites located in the 3’UTR of target mRNAs has been the focus of the majority of miR-based studies. However, in depth analysis by Fang & Rajewsky (66) and our own recent characterization of miR targets (42,61) have shown that miR target sites in the CRS are of functional importance, are under negative selection and that target sites in the CRS can enhance regulation mediated by sites in the 3’UTR. Based on these findings it is likely that the two miR-128 binding site in TNPO3 mRNA corporate for optimal repression of TNPO3 expression.

The finding that TNPO3 is a direct target of miR-128 could indicate that miR-128 functions by interfering with TNPO3/CPSF6-dependent nuclear import of HIV-1 and viral replication. However, both infection studies using wild-type and N74D mutant HIV-1 reporter virus and CPSF6-depletion studies (in which the effect of CPSF6 removal was modest) suggest that there are other miR-128 targets that can also regulate HIV-1 infection. Our work does suggest that miR-128 functions in a TNPO3-dependent and TNPO3-independent context, during HIV-1 infection. However, further mechanistic studies are needed to evaluate whether miR-128 directly blocks nuclear import of HIV-1 PIC and exactly where in the viral life cycle the anti-viral effect of miR-128 can be observed.

As mentioned we have previously established a novel role for miR-128 in the inhibition of long-interspaced elements-1 (LINE-1 or L1) retrotransposition (42). Interestingly, miR-128-induced L1 restriction takes place by both direct interactions with L1 RNA and indirectly via miR-128-induced depletion of cell host factors, which L1 is dependent on for successful replication (42 and *manuscript under revision*). From an evolutionary standpoint miRs have likely evolved to protect us against unwanted L1 mobilization, including cancer-initiating mutagenesis, which could lead to detrimental genomic instability. The network of cellular host factors and pathways which mobile DNA elements are dependent on, also restrict the life cycle of retrovirus, and as such it seems likely that miRs (including miR-128) play a role in the potent arsenal of anti-viral defense factors in human cells, also acting against HIV-1 infection. As miRs are known to often function by regulating multiple gene products in the same cellular pathway (67-70), we predict that miR-128 regulates multiple cellular co-factors some of which HIV-1 is dependent on and as such miR-128 may function as a master restriction factor of RNAs stemming from extracellular pathogens including HIV-1, in addition to intracellular selfish elements such as L1.

The finding that miR-128 restricts L1 replication by a dual mechanism, by regulating cellular cofactors (including TNPO1) and by targeting L1 RNA, raises the question as to whether miR-128 may also directly target the HIV-1 genome. However unlike L1, which is not capable of responding to selective pressure, HIV-1 would be predicted to evolve by mutating the miR-128 binding site(s) and escape miR-128-induced inhibition, *if* miR-128 is potently restricting viral replication.

In summary, our results support the model that miR-128 which is expressed in primary HIV-1 target cells and is a type I IFN response gene, functions as a novel anti-viral defense mechanism during HIV-1 infection, partly by repressing the nuclear import factors TNPO3 and inhibiting HIV-1 replication (see Figure 6). A long-term goal of our mechanistic analysis including network studies is to add to the understanding of the interactome of viral infection, replication, latency and reactivation which will enable us to propose novel therapeutic strategies to target specific co-factors that could prevent successful HIV-1 infection, block viral replication or attenuate establishment/reactivation of latency, all components of a functional cure.

**Figure 6:**
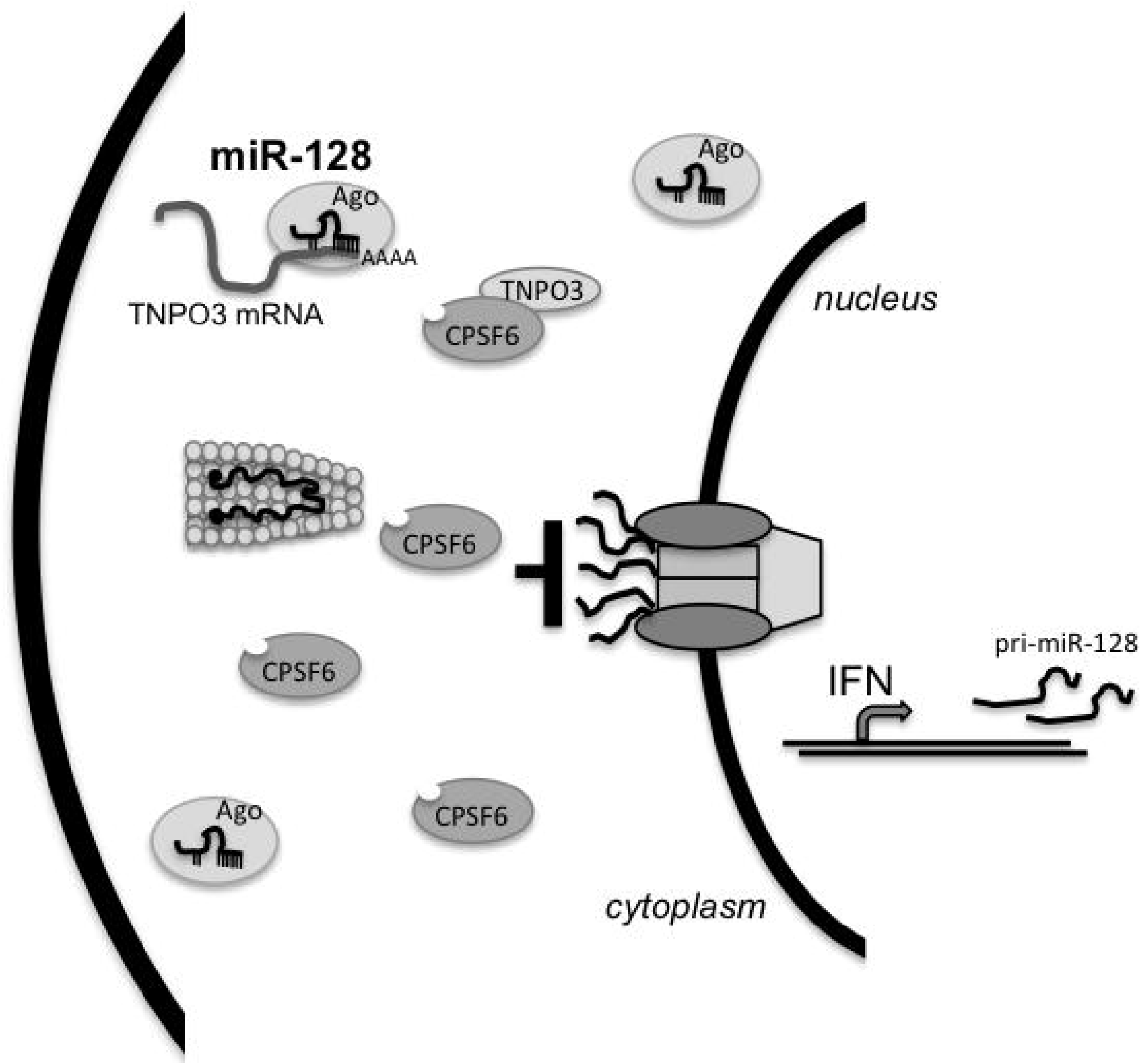
Model of miR-128-induced inhibition of HIV-1 replication by targeting TNPO3. miR-128 is induced by type I interferon and directly targets TNPO3 mRNA. Loss of TNPO3 results in accumulation of CPSF6 in the cytoplasm and inhibition of viral nuclear import and HIV-1 replication.

## ACKNOWLEDGEMENTS

This work was supported by University of California Cancer Research Coordinating Committee 55205 (I.M.P.), American Cancer Society – Institutional Research Grant 98-279-08 (I.M.P.), University of California Irvine Institute for Memory Impairments and Neurological Disorders grant (I.M.P.).

## AUTHOR CONTRIBUTIONS

A. Bochnakien performed the majority of experiments, with the help of A. Idica and M Hamdorf, demonstrating that miR-128 targets TNPO3 and that miR-128 specifically inhibit HIV-1 replication, using single cycle reporter virus. D. Zisoulis performed the Ago RNA IPs. K. Lee and A. Zhen performed spreading experiments in cell lines and in primary CD4 T-cells. V. KewalRamani and S. Kitchen contributed expertise and advise on infection and HIV biology experiments. I. Daugaard generated the final figures and commented on the manuscript. IM. Pedersen performed some experiments and directed all other experiments, figure design and wrote the manuscript. All authors reviewed the results and approved the final version of the manuscript.

## CONFLICT OF INTEREST

The authors declare no conflict of interest.

## MATERAILS AND METHODS

### Cell culture and primary HIV-1 target cell isolation

All cells were cultured at 37°C and 5% CO2. HeLa cells (CCL-2), 293T cells (CRL-3216), THP-1 cells (TIB-202) and Jurkat cells (TIB-152) were all obtained from ATCC. Adherent cells were cultured in EMEM (SH3024401, Hyclone) supplemented with 10% HI-FBS (FB-02, Omega Scientific), 5% Glutamax (35050-061, ThermoFisher), 3% HEPES (15630-080, ThermoFisher). Jurkat cells were cultured in RPMI-1640 medium supplemented with 10% HI-FBS (FB-02, Omega Scientific).

Peripheral blood mononuclear cells (PBMC) were isolated from whole blood obtained from anonymous blood donors (New York Blood Center) using a standard Ficoll (Cellgro) procedure. Primary CD4 T cells isolated using the human CD4 T cell enrichment kit per the manufacturer’s instruction (Stemcell) were activated using Dynabeads Human T-Activator CD3/CD28 (Life Technologies) and cultured in suspension cell medium supplemented with 30 units per ml of interleukin-2 (IL-2) (Pepro-Tech).

Alternatively primary human blood derived monocytes were isolated from peripheral blood mononuclear cells after Ficoll density gradient centrifugation. PBMC were allowed to attach to a 10-cm dish for 2 h and then vigorously washed with phosphate-buffered saline (PBS). Attached cells were subjected to differentiation in RPMI supplemented with 10% FBS, P/S, 2 mM2-glutamine, and 100 ng per ml of granulocyte macrophage colony-stimulating factor (GM-CSF) (PeproTech) for 4 days. Some primary CD4 T cell and primary blood-derived monocyte cultures were stimulated with interferon-alpha (100U/mL) (PeproTech) for 72-96 hours.

### Viral Infections

All HIV-1 reporter vectors encoding fluorescent proteins or firefly luciferase were NL4-3 derived. Reporter particles incorporating VSV-G and replication competent HIV-1 were derived through transient transfection of HEK293T cells. For some cultures an MLV-RFP/VSV-G reporter was used to evaluate infection specificity, as compared to HIV-RFP/VSV-G. The percentage of infected cells were measured by FACS analysis of RFP expression.

Alternatively, stable miR modulated Jurkat cells were spinfected with wildtype HIV-1_(NL4-3)_ virus and viral replication was determined by p24 ELISA assays after 7, 9 and days of infection. Equal infection was confirmed by normalizing to RT units or virus was pre-titered on GHOST cells and equal infection units were used.

Finally, activated primary CD4 T-cells were infected with HIV-1 NL4-3 (100ng/10E6) for 7 days. Supernatant were collected on day 3,5 and 7 for p24 assays. On day 7 infected cells were harvested and stained for intracellular anti-gag (clone KC57). Level of infection as presented by % gag+ is shown among GFP+ transduced cells.

### Transfection and transduction

OptiMem (31985070, Lifetech) and Lipofectamine RNAiMAX (13778075, Lifetech) were used to complex and transfect 20μM miR-128 mimic, anti-miR-128 or control mimics (C-301072-01 and IH-301072-02, Dharmacon) into cells. OptiMem and Xtreme Gene HP (06366236001, Roche Lifescience) was used to transfect pJM101 neomycin L1 reporter plasmid into HeLa cells. Cells were transduced with high titer virus using polybrene (sc-134220, Santa Cruz Biotech) and spinoculation (800xg at 32°C for 30 minutes). Transduced cells were then selected and maintained using 3μg/mL puromycin.

### RNAi using shRNA against TNPO3

shRNA for TNPO3 was designed using the RNAi Consortium (https://www.broadinstitute.org/rnai/public/) using clone TRCN0000235098 and cloned into pLKO.1 puro backbone (Addgene, #8453). pLKO shGFP control plasmid was pre-assembled (Addgene, #30323).

### Lentiviral packaging

VSVG-pseudotyped lentiviral vectors were made by transfecting 0.67μg of pMD2-G (12259, Addgene), 1.297μg of pCMV-DR8.74 (8455, Addgene), and 2μg of mZIP-miR-128, mZIP-anti-miR-128, pLKO-shControl or pLKO-shHNRNPA1 (transfer plasmid)) into 293T cells using Lipofectamine LTX with plus reagent (15338030, ThermoFisher). Virus-containing supernatant was collected 48hr and 96hr post-transfection. Viral SUPs were concentrated using PEG-it virus precipitation solution (LV810A-1) according to manufacturer’s instructions.

### RNA extraction and quantification

RNA was extracted using Trizol (15596-018, ThermoFisher) and Direct-zol RNA isolation kit (R2070, Zymo Research). cDNA was made with High-Capacity cDNA Reverse Transcription Kit (4368813, ThermoFisher). Amount of TNPO3 mRNA was analyzed by qRT-PCR (Sense primer 5’-aagcaattttggaggtggtg-3’; Antisense primer 5’-atagccaccttggtttcgtg-3’) using Forget-me-not qPCR mastermix (Biotium) relative to beta-2-microglobulin (B2m, Sense primer 5’-ATGTCTCGCTCCGTGGCCTTAGCT-3’; Antisense primer 5’-TGGTTCACACGGCAGGCATACTCAT-3’) housekeeping gene and processed using the ΔΔC_t_ method.

### Western blotting

Rabbit anti-TNPO3 antibody (ab109386, Abcam) was used at 1:2000. Rabbit anti-CPSF6 antibody (ab99347, Abcam), Anti-alpha-Tubulin antibody (ab4074, Abcam) was diluted 1:5000 and used as a loading control, validation of antibodies can be found on the manufacturer websites. Secondary HRP-conjugated anti-rat (ab102172, Abcam) or HRP-conjugated anti-rabbit (#NA934, GE lifesciences) were used at 1:5000. ECL substrate (32106, ThermoFisher) was added and visualized on a BioRad ChemiDoc imager.

### Argonaute (Ago)-RNA immuno-purification

Immunopurification of Argonaute from HeLa cell extracts was performed using the 4F9 antibody (#sc-53521, Santa Cruz Biotechnology) as described previously (42). Briefly, 10mm plates of 80% confluent cultured cells were washed with buffer A [20 mM Tris-HCl pH 8.0, 140 mM KCl and 5 mM EDTA] and lysed in 200ul of buffer 2XB [40 mM Tris-HCl pH 8.0, 280 mM KCl, 10 mM EDTA, 1% NP-40, 0.2% Deoxycholate, 2X Halt protease inhibitor cocktail (Pierce), 200 U/ml RNaseout (Life Technologies) and 1 mM DTT. Protein concentration was adjusted across samples with buffer B [20 mM Tris-HCl pH 8.0, 140 mM KCl, 5 mM EDTA pH 8.0, 0.5% NP-40, 0.1% deoxycholate, 100 U/ml Rnaseout (Life Technologies), 1 mM DTT and 1X Halt protease inhibitor cocktail (Pierce)]. Lysates were centrifuged at 16,000g for 15 mins at 4^o^C and supernatants were incubated with 10-20 ug of 4F9 antibody conjugated to epoxy magnetic beads (M-270 Dynalbeads, Life Technologies) for 2 hours at 4^o^C with gentle rotation (Nutator). The beads, following magnetic separation, were washed three times five mins with 2 ml of buffer C [20 mM Tris-HCl pH 8.0, 140 mM KCl, 5 mM EDTA pH 8.0, 40 U/ml Rnaseout (Life Technologies), 1 mM DTT and 1X Halt protease inhibitor cocktail (Pierce)]. Following immunopurification, RNA was extracted using miRNeasy kits (QIAGEN), following the manufacturer’s recommendations and qPCR was performed using hn RNPA1 primers designed around the binding site of miR-128 (Sense primer 5’-TCTCCTAAAGAGCCCGAACA-3’; Antisense primer 5’-TTGCATTCATAGCTGCATCC-3’) or GAPDH (Sense primer 5’-GGTGGTCTCCTCTGACTTCAA-3’; Antisense primer 5’-GTTGCTGTAGCCAAATTCGTT-3’) normalized to B2m (Sense primer 5’-ATGTCTCGCTCCGTGGCCTTAGCT-3’; Antisense primer 5’-TGGTTCACACGGCAGGCATACTCAT-3’). Results were normalized to their inputs.

### Luciferase binding assay

Wild-type TNPO3 sense primer (5’-AATTCTTGGGTTTGTCACATATGCCACTGTGGAGGAGGTGGATGCAGCTA-3’) and antisense primer (5’-CTAGTAGCTGCATCCACCTCCTCCACAGTGGCATATGTGACAAACCCAA-3’), mutated TNPO3 sense primer (5’-AATTCTTGGGTTTGTCACATATGCCCTTATGGAGGAGGTGGATGCAGCTA-3’) and antisense primer (5’-CTAGTAGCTGCATCCACCTCCTCCATAAGGGCATATGTGACAAACCCAA-3’), or positive control sense primer (5’-AATTCAAAGAGACCGGTTCACTGTGAA-3’) and antisense primer (5’-CTAGTTCACAGTGAACCGGTCTCTTTG-3’) sequences were cloned into dual-luciferase reporter plasmid (pEZX-MT05, Genecopoeia). 3x10^5^ HeLa cells were forward transfected with 0.8μg of reporter plasmid (WT, mutated, Pos) and 20nM miR-128 mimic (Dharmacon) or Control mimic (Dharmacon) using Attractene transfection reagent (301005, Qiagen) according to manufacturer instructions. Relative Gaussia Luciferase and SEAP was determined using Secrete-Pair Dual Luminescence Assay Kit (SPDA-D010, Genecopoeia). Luminescence was detected by Tecan Infinite F200 Pro microplate reader.

### Statistical analysis

Student’s t-tests were used to calculate two-tailed *p* values and data are displayed as mean ± standard error of the mean (SEM) of technical (TR) or independent biological replicates (IBR), (n) as indicated.

---

**Figure S1:**
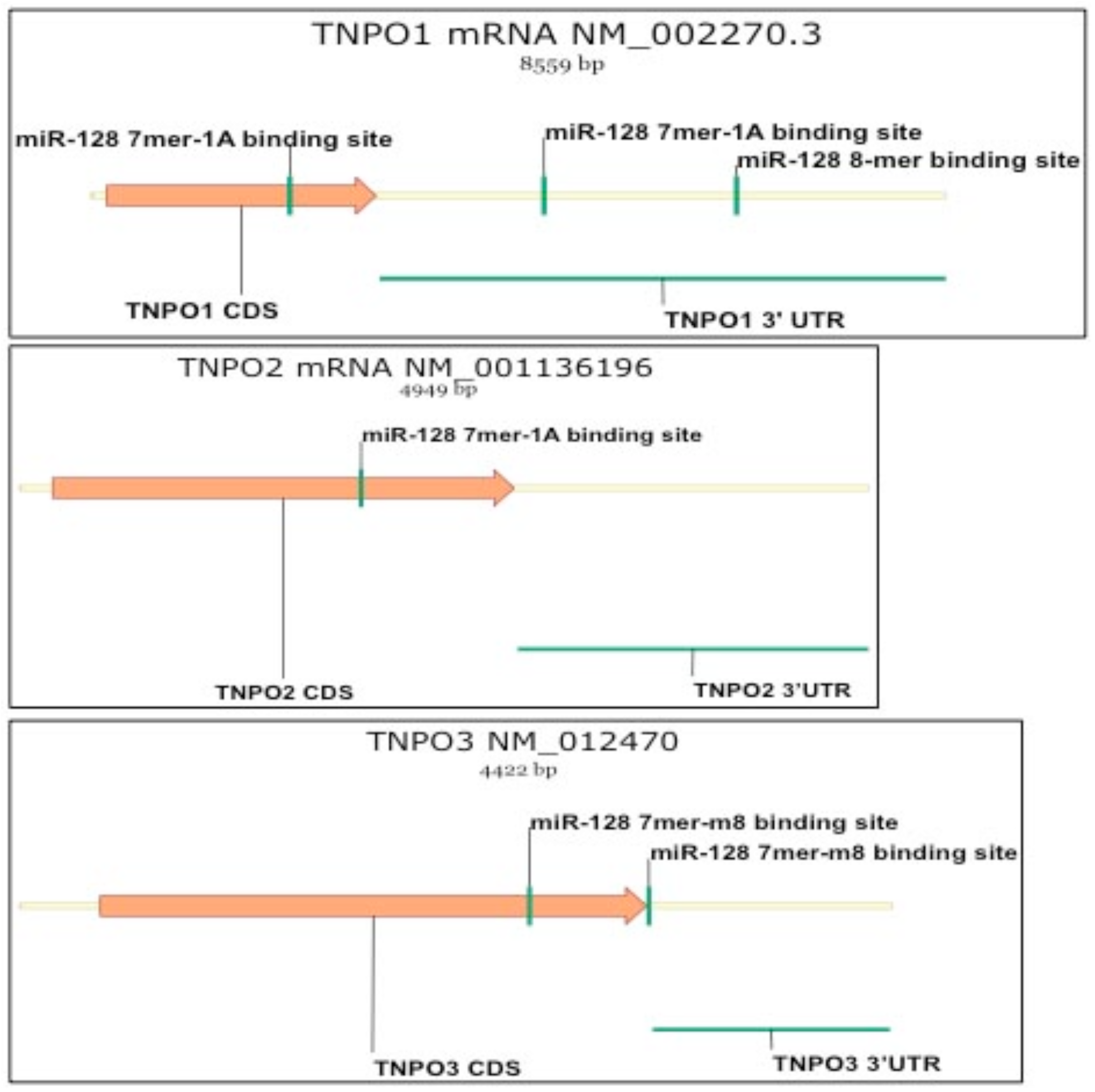
Predicted miR-128 binding sites in the three different TNPO isoforms (TNPOl, TNP02 and TNP03).

**Figure S2:**
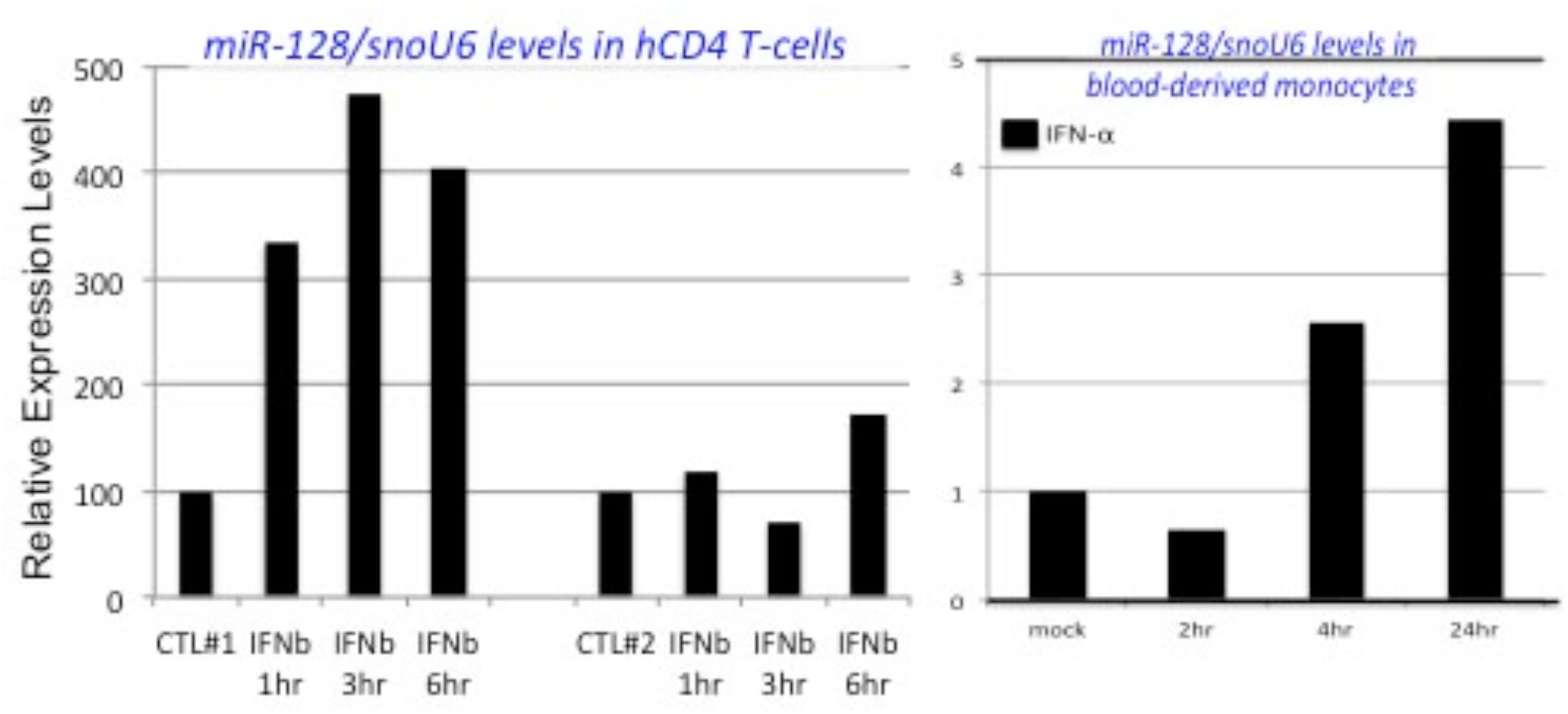
miR-128 expression and induction by type 1 interferon. Primary human CD4 T-cells and blood derived monocytes were isolated from healthy donors by Ficoll and MACS separation versus adhesion separation. Cells were cultured in the presence or absence of type 1 interferon (lOOU/mL). Cells were harvested and miR specific qPCR was performed for miR-128, normalized to snoU6.

**Figure S3:**
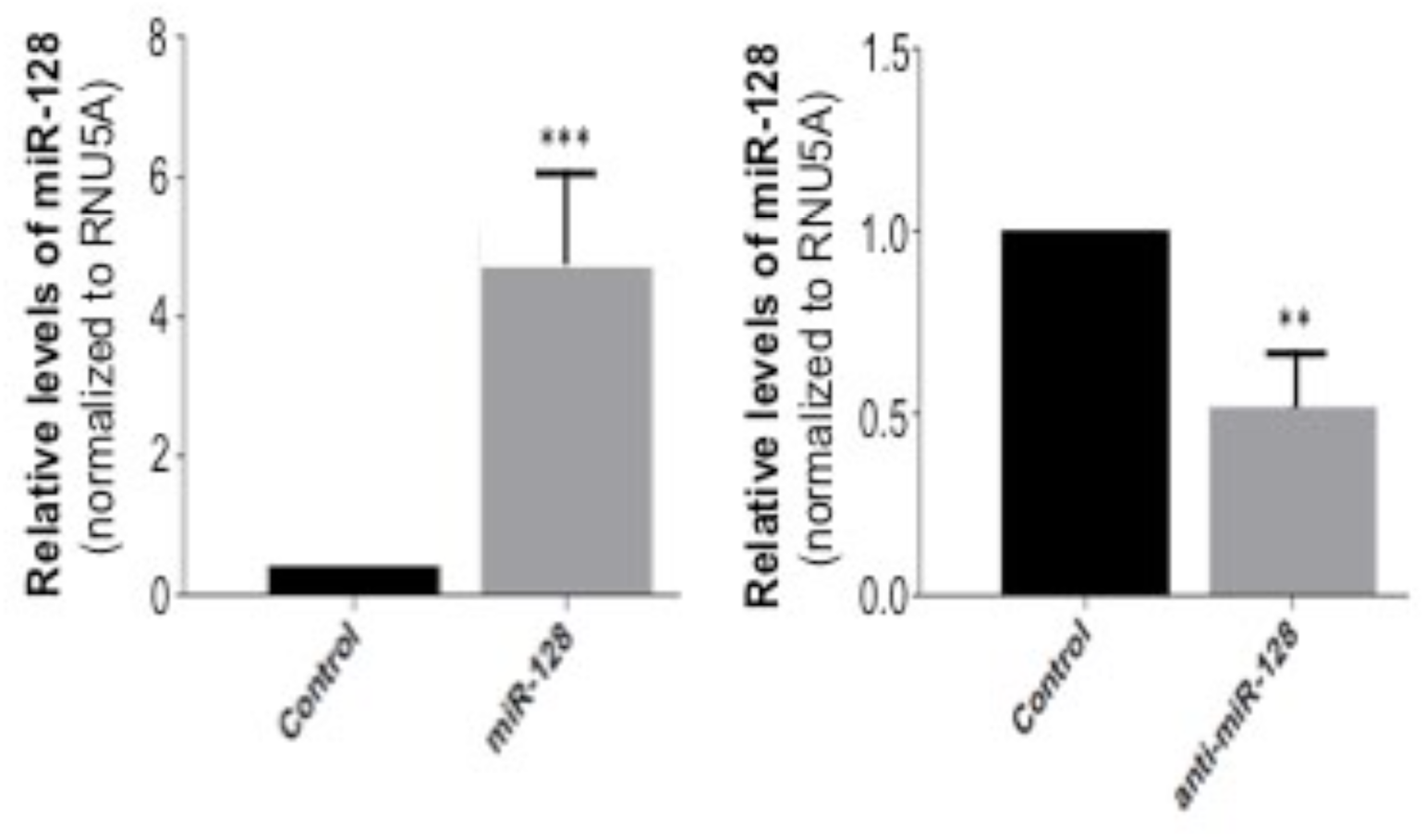
miR-128 expression verification in miR modulation HeLa cells. miR specific qPCR was performed on stable miR modulated HeLa cells using miR-128 specific primers, and normalizing to RNU5A.

**Figure S4:**
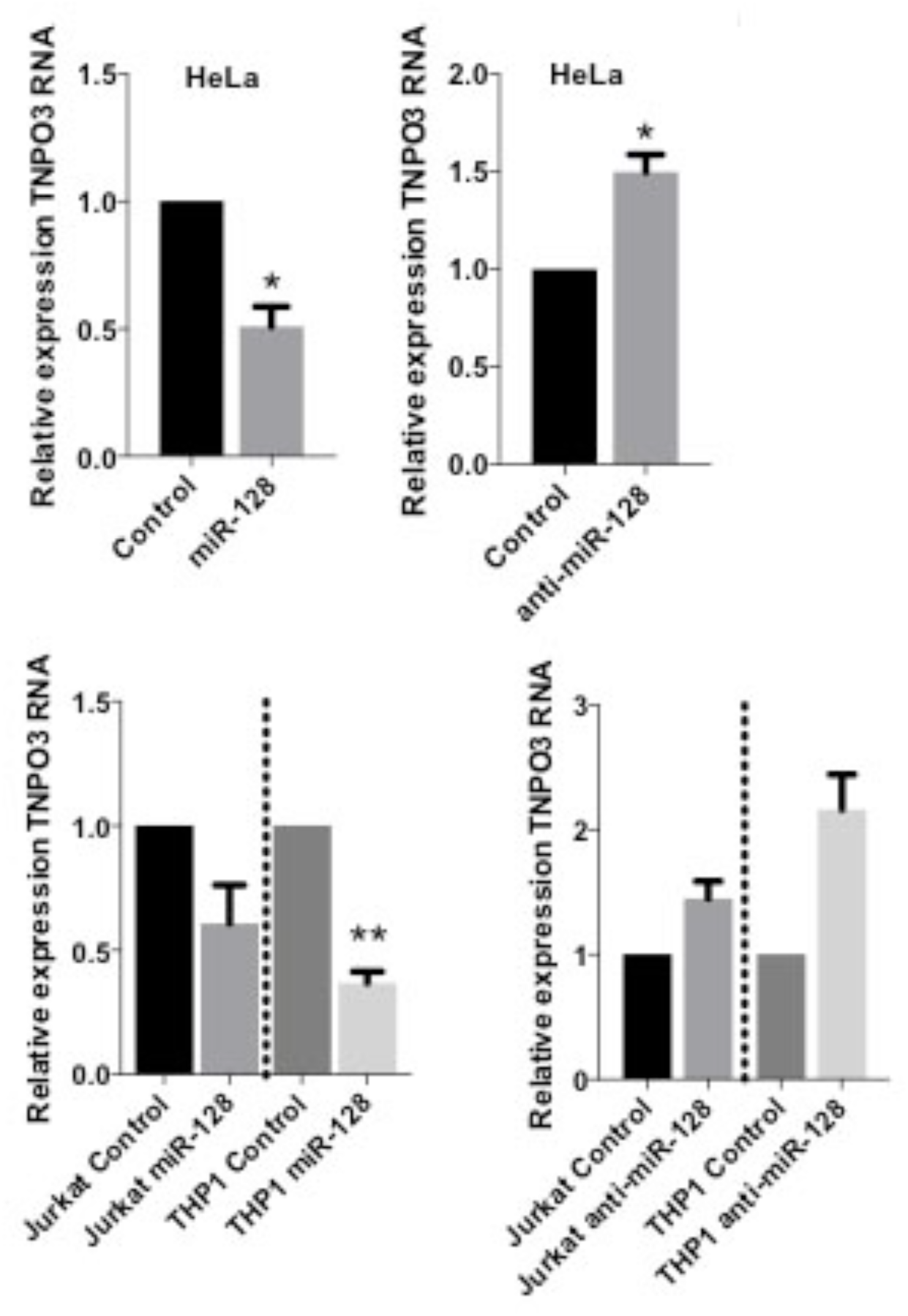
miR-128 regulation of TNP03 mRNA by transient transfections. (A) Relative levels of TNP03 RNA to B2M in HeLa, Jurkat or THP-1 cells transiently transfected with control miR, anti-miR-128 or miR-128 are shown as mean t SEM (n=3, independent biological replicate) Throughout figure, *P<0.05; **P<0.01 ***, by two-tailed Student’s *t* test.

